# Large differences in small RNA composition between human biofluids

**DOI:** 10.1101/251496

**Authors:** Paula M. Godoy, Nirav R. Bhakta, Andrea J. Barczak, Hakan Cakmak, Susan Fisher, Tippi C. Mackenzie, Tushar Patel, Richard W. Price, James F. Smith, Prescott G. Woodruff, David J. Erle

## Abstract

Extracellular miRNAs and other small RNAs are implicated in cellular communication and may be useful as disease biomarkers. We systematically compared small RNAs in 12 human biofluid types using RNA-seq. miRNAs and tRNA-derived RNAs (tDRs) accounted for the majority of mapped reads in all biofluids, but the ratio of miRNA to tDR reads varied from 72 in plasma to 0.004 in bile. miRNA levels were highly correlated across all biofluids but levels of some miRNAs differed markedly between biofluids. tDR populations differed extensively between biofluids. Y RNA fragments were seen in all biofluids and accounted for >10% of reads in blood plasma, serum, and CSF. Reads mapping exclusively to piRNAs were very rare except in seminal plasma. These results demonstrate extensive differences in small RNAs between human biofluids and provide a useful resource for investigating extracellular RNA biology and developing biomarkers.

## INTRODUCTION

RNAs released from cells have been detected in many biofluids (Patton et al., 2015). Although some reports suggest that large RNAs, including functional mRNAs, can be present in biofluids, (Ni et al., 2002; Skog et al., 2008), most extracellular RNAs (exRNAs) are small RNAs (Hoy and Buck, 2012). Many reports have focused on miRNAs, which are small (typically 21-22 nt) RNAs produced by intracellular processing of larger precursor RNAs (Argyropoulos et al., 2013; Lee et al., 2010; Mitchell et al., 2008; Weber et al., 2010; Williams et al., 2013). Several pathways have been proposed to lead to release of miRNAs from cells and extracellular miRNAs can be found within exosomes and in complexes containing argonaute proteins (Arroyo et al., 2011) or lipoproteins (Patton et al., 2015). Extracellular miRNAs can enter cells and may target mRNAs in those cells (Patton et al., 2015). The potential of miRNAs as disease biomarkers is being explored by many investigators (Argyropoulos et al., 2013; Barger et al., 2016; Gray et al., 2017; Mitchell et al., 2008).

Small RNA biotypes other than miRNAs have also been detected in biofluids. Piwi-interacting RNAs (piRNAs) are ∼23-30 nt RNAs involved in transcriptional and post-transcriptional silencing of transposons and other targets in germ cells (Czech and Hannon, 2016). Sequences mapping to piRNAs have been reported in human seminal fluid plasma (Hong et al., 2016) as well as in blood plasma (Freedman et al., 2016), saliva (Bahn et al., 2015) and urine (Yeri et al., 2017). Many small RNAs found in biofluids are thought to be derived from larger RNAs by specific processing events or by non-specific degradation. Transfer RNA-derived RNAs (tDRs) are generated by cleavage of tRNAs at specific sites (Gebetsberger and Polacek, 2013). The variety of tDRs is extensive since there are many (>600) human tRNA genes and multiple fragment types, including 5’-halves, 3’-halves, 5’-tRNA-fragments (tRFs), 3’-tRFs, and internal tRFs (i-tRFs) have been identified (Loher et al., 2017). Small RNAs derived from mRNAs, long non-coding RNAs, ribosomal RNAs (Semenov et al., 2004), and Y RNAs (Dhahbi et al., 2013) have also been detected in some biofluids. Their functions are not yet well understood.

Extracellular RNAs have been detected in at least 15 biofluids to date (Sohel, 2016). The total concentration of RNA varies widely between biofluids, with certain biofluids such as breast milk and seminal fluid being more concentrated than more dilute biofluids like CSF and urine (Weber et al., 2010). Given the difficulties inherent in absolute quantification of exRNAs, most analyses have focused on relative quantification of specific RNAs. This approach has identified some differences in small RNA content between biofluid types. For example, a PCR-based study of 714 miRNAs identified certain miRNAs that were abundant in most of the 12 biofluids studied along with other miRNAs that were enriched in specific biofluids (Weber et al., 2010). However, a more complete understanding of small RNA differences between biofluids is problematic since published studies rely on a diverse set of methods with different biases that preclude direct comparisons. Furthermore, most studies have focused primarily or exclusively on miRNAs and much less information is available about other RNA biotypes.

One objective of the National Institutes of Health Extracellular RNA Communication Consortium (ERCC) is to identify the range of RNAs present in human biofluids. To address this goal, we compared the small RNA populations of a large and diverse collection of human biofluids using one standard RNA-seq approach that was validated as part of a multicenter ERCC study (Giraldez et al., 2017). This approach was designed for miRNAs but can also detect other small RNAs with a 5’ phosphate and a 3’ hydroxyl group. Our analysis of a total of 129 samples of 12 biofluid types from human donors reveals the presence of complex RNA repertoires in all biofluids and major differences in RNA composition between biofluid types. These results are publicly available through the exRNA Atlas (https://exrna-atlas.orgx).

## RESULTS

### Characteristics of the study population

We obtained samples of 12 different biofluid types (Table 1). With the exception of bile samples, which were obtained at the Mayo Clinic from patients who had previously undergone liver transplantation and had intact liver function, all samples were obtained from healthy subjects in the course of studies performed at UCSF. For each biofluid, we obtained 5-15 samples. In general, each sample of a given biofluid was obtained from a different participant, except that the 10 ovarian follicle fluid samples were obtained from five participants who each provided two samples. Plasma, serum, and urine samples were obtained from a separate cohort of 12 individuals who each provided samples of each of these three biofluids. For saliva, another cohort of 15 participants provided samples of both parotid saliva and submandibular/sublingual (SMSL) saliva. For amniotic fluid, BAL fluid, bile, cord blood plasma, CSF, seminal fluid, 10-12 participants from separate cohorts each provided a single sample of one biofluid.

**Table 1.** Study Participant Demographic Summary.

| Biofluid | Sex<sup>a</sup> | Age (years)<sup>b</sup> | Race |  |  |  |  |  | Ethnicity |  |  |  |
| --- | --- | --- | --- | --- | --- | --- | --- | --- | --- | --- | --- | --- |
|  |  |  | African American | Asian | White | Other (mixed) | Unknown | Pacific Islander | Hispanic | Non-Hispanic | Declined to State | Unknown |
| Amniotic Fluid | 10F | Unknown |  |  |  |  | 10 |  |  |  |  | 10 |
| BAL fluid | 4F, 6M | 29.5 (28.3-33.0) | 3 | 1 | 6 |  |  |  | 1 | 9 |  |  |
| Bile | 6F, 6M | 57.5 (53.3-60.3) |  |  | 12 |  |  |  |  | 11 | 1 |  |
| Cord Blood Plasma (Maternal demographics) | 10F | 31 (29.5-33.5) | 1 |  | 4 | 2 | 2 |  | 3 | 5 |  | 2 |
| Cord Blood Plasma (Fetal demographics) | 6F, 4M | 0 (0-0) | 1 | 2 | 2 | 3 | 2 |  | 3 | 4 |  | 3 |
| CSF | 3F, 8M | 40 (37.5-43.0) | 6 | 1 | 4 |  |  |  |  | 11 |  |  |
| Ovarian Follicle Fluid | 5F | 28 (26-28) |  | 2 | 3 |  |  |  |  | 5 |  |  |
| Plasma | 9F, 3M | 26 (24.8-35.3) |  | 7 | 4 |  |  | 1 | 2 | 10 |  |  |
| Serum | 8F, 3M | 26 (24.5-30.0) |  | 7 | 3 |  |  | 1 | 1 | 10 |  |  |
| Urine | 8F, 2M | 27.5 (25.3-43.8) |  | 7 | 2 |  |  |  | 1 | 9 |  |  |
| Parotid Saliva | 8F, 5M | 29 (24-30) |  | 3 | 8 |  | 2 |  | 2 |  |  | 11 |
| SMSL Saliva | 8F, 7M | 29 (27-33) |  | 3 | 10 |  | 2 |  | 2 |  |  | 13 |
| Seminal Fluid | 10M | 40.5 (33.8-40.5) | 1 | 3 | 6 |  |  |  | 1 | 9 |  |  |

<sup>a</sup> F, female; M, male
<sup>b</sup> Median (interquartile range)

### Sequencing and quality control

RNA-seq reads were aligned to the human transcriptome using the Genboree/exceRpt pipeline (http://www.genboree.org). By default, the pipeline assigns reads in the order miRNA, tRNA, piRNA, “Gencode” (other Ensembl transcripts including mRNAs and non-coding RNAs including Y RNAs, small nuclear RNAs, and long non-coding RNAs), and then circular RNA. This approach resulted in a substantial proportion of read counts assigned to piRNAs in adult blood plasma (median 13% assigned to piRNA), cord blood plasma (5%), and serum (6%) samples. Very large proportions of reads that mapped to piRNA in these three biofluids mapped to a single piRNA sequence (piRNAbase hsa_piR_016658, Genbank identifier DQ592931; 79% of piRNA reads in adult blood plasma, 48% in cord blood plasma, and 43% in serum). However, these reads also map to a Y RNA (RNY4) and each of these biofluids also contain large numbers of other reads that map to other regions of RNY4. In the aggregate, >99.7% of plasma or serum reads mapping to piRNAs also mapped to RNA(s) from a different biotype. Based upon these observations, we changed the order of read assignment to miRNA, tRNA, Gencode, piRNA, and then circular RNA. Using this approach, 0.3% of seminal plasma reads mapped to piRNAs whereas <0.1% of reads in other biofluids mapped to piRNAs.

A total of 145 samples were analyzed. 16 samples failed quality control despite repeat analysis (see Experimental Procedures). The remaining 129 samples were used for the analyses presented here. The median number of reads aligning to the human transcriptome was 3.6 ✕ 10^6^ (interquartile range [IQR]: 0.8-8.3 ✕ 10^6^).

### Relative abundances of RNA biotypes differ widely between biofluids

The fraction of reads mapping to different biotypes of human RNAs varied markedly between biofluid types (Fig. 1, Table S1). The median rate of mapping to miRNAs was >50% for adult and cord blood plasma and BAL fluid, whereas tDRs represented >50% of reads in bile, urine, seminal plasma, and amniotic fluid. Both miRNAs and tDRs each accounted for >25% of reads in the remaining biofluids (parotid saliva and SMSL saliva, ovarian follicle fluid, serum, and CSF). The relative abundance of miRNA to tRNA reads varied by >10^3^-fold (from 72 in plasma to 0.004 in bile). Y RNA fragments represented >10% of mapped reads in adult and cord blood plasma, serum, and CSF but <0.8% of reads in urine and bile, with intermediate levels in other biofluids. miRNAs, tRNAs, and Y RNAs accounted for >90% of all mapped reads except in SMSL saliva, which had the largest proportion of reads mapping to portions of protein coding genes (mRNAs, 7.6%), snRNAs (2.3%), retained introns (1.5%), and Gencode “processed transcripts” (1.2%). This may reflect the presence of RNA degrading enzymes, cellular debris, or microbial-derived small RNAs that map to the human genome in saliva samples. Reads that mapped to piRNAs but not to miRNAs, tRNAs, Y RNAs, or other Gencode transcripts represented 0.30% of reads in seminal plasma and <0.10% of reads in other biofluids. These results demonstrate that multiple classes of small RNAs are represented in each biofluid type studied, but the relative abundance of these classes varies widely between biofluids.

**Figure 1.**
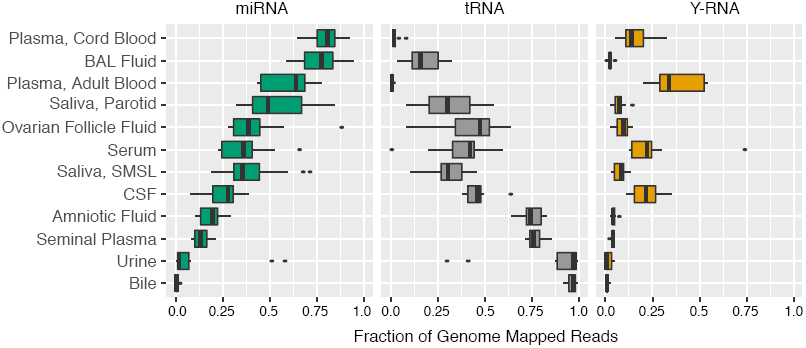
Distribution of RNA biotypes differs between biofluids. Reads mapping to miRNAs, tRNAs, Y-RNAs, piRNAs, mRNAs, or other RNA biotypes as a fraction of total reads mapping to the human transcriptome. Boxes represent median and interquartile ranges, whiskers represent 1.5 times the interquartile range. Dots represent outliers.

### miRNA profiles

In each biofluid, we detected hundreds of miRNAs but a small number of miRNAs accounted for a large proportion of miRNA read counts (Fig. 2A-B, Fig. S1, and Table S2). Between 395 (parotid saliva) and 541 (CSF) miRNAs had a median of ≥10 reads/million total miRNA reads in each biofluid. The 10 most frequent miRNAs represented between 39% (CSF) and 62% (serum) of total miRNA reads (Table S3). Despite the marked differences in small RNA classes between biofluids, pairwise correlation coefficients for miRNA read counts between biofluids were generally high (*R* = 0.79 − 0.98, Fig. 2C-E). Correlations were highest between blood-derived biofluids (plasma, cord plasma, and serum; *R* = 0.94-0.98) and between saliva samples from the two different sites (*R* = 0.98). Seminal fluid and cord blood plasma were the least correlated (*R* = 0.79). We used a dimensionality reduction method, transfer stochastic neighbor embedding (tSNE), to determine whether there were consistent differences in the small miRNA composition of the 12 biofluid types. Using all 2,153 miRNAs detected in any sample, tSNE analysis revealed that samples of most biofluid types formed distinct clusters (Fig. 2F). Samples of saliva from two different sites (parotid and submandibular/sublingual glands) formed overlapping clusters and two serum samples could not be clearly distinguished from the cluster of adult plasma samples. Therefore, although correlations in miRNA levels between biofluids were high, each biofluid (with the exception of the saliva samples from two different sites) had a distinct pattern of miRNAs.

**Figure 2.**
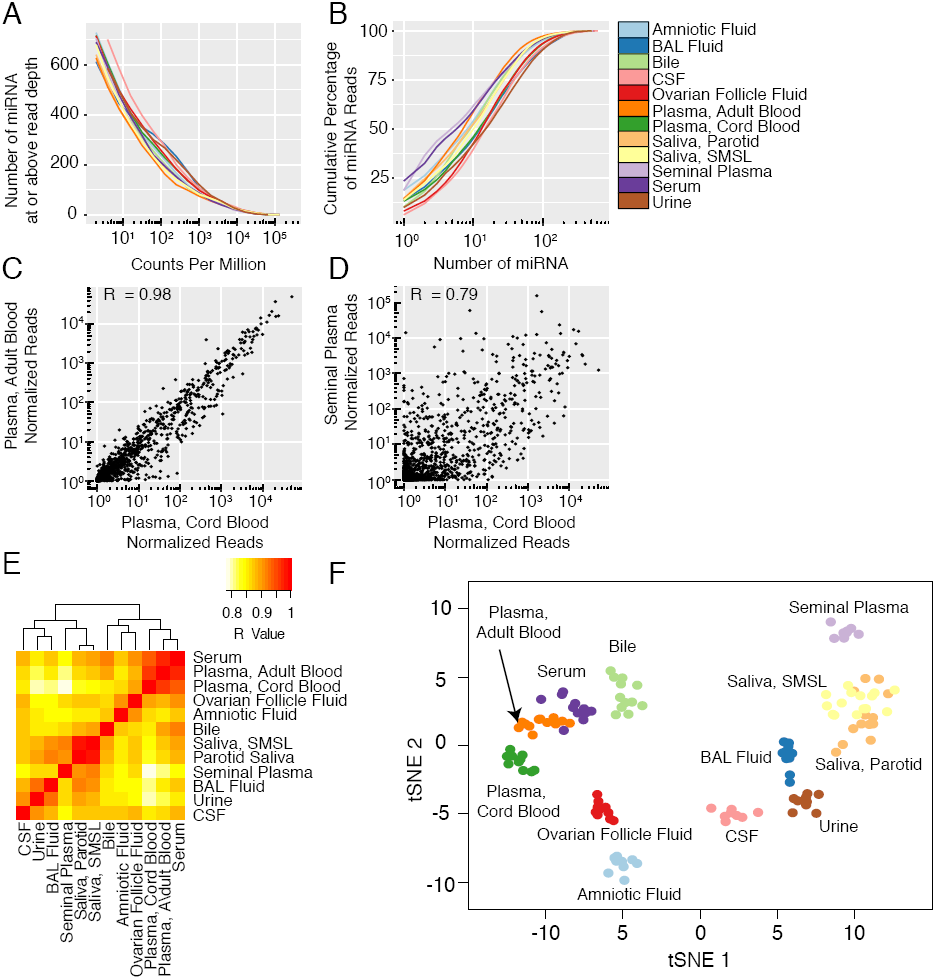
miRNA profiles in 12 biofluid types. (A) Number of miRNAs detected as a function of read depth. (B) Cumulative distribution of miRNA reads. (C and D) Examples of pairwise correlations between biofluids. Each point represents the median normalized read count for a single miRNA for the indicated biofluids. One normalized read count was added to each measurement to allow representation of log read counts for miRNAs with no reads. (E) Correlations for all pairs of biofluids. (F) tSNE plot produced using miRNA read counts. Each point represents a single biofluid sample.

A total of 15 miRNAs had much higher relative abundance in one biofluid than any other biofluid (≥10^3^ reads/10^6^ total miRNA reads, >10-fold higher in one biofluid than all other biofluids, adjusted p<0.05 for pairwise comparisons with all other biofluids by negative binomial Wald Test, Table S4). In some cases, these levels are likely explained by high expression of the miRNAs by cell types that are in direct contact with the biofluid. Three of the four miRNAs with higher levels in amniotic fluid (miR-483-5p, miR-1247-5p, and miR-433-3p) are highly enriched in extraembryonic cells (amniotic epithelial cells, placental epithelial cells, or chorionic membranes) (de Rie et al., 2017). The three miRNAs with much higher levels in CSF are miR-9-3p, which is highly enriched in the vertebrate nervous system; miR-1911-5p, a brain-specific miRNA that was detected in CSF exosomes but not blood plasma exosomes (Yagi et al., 2017), and miR-1298-5p, which is among the most abundant miRNAs in CSF exosomes (Yagi et al., 2017). miR-891a, which had much higher expression in seminal plasma, has been reported to be among the most abundant miRNAs in epididymis (Li et al., 2012). Combining results all blood-derived biofluids (adult and cord blood plasma and serum) into a single group and comparing with results each of the other biofluids did not identify any miRNAs that met the above criteria for much higher expression in blood-derived biofluids versus all other biofluids. Similarly, combining results from parotid and SMSL saliva samples did not identify any miRNAs with much higher expression in saliva versus all other biofluids.

A comparison of umbilical cord blood plasma and adult blood plasma revealed that 18 miRNAs differed in relative abundance (Table S5). Three miRNAs, miR-487b-3p, miR-376c-3p, and miR-127-3p were >5-fold higher in cord blood plasma. A previous report detected similar differences between cord and adult blood plasma and showed relatively high levels of each of these miRNAs in placenta (Williams et al., 2013). One miRNA, let-7b-5p, was >10-fold lower in cord blood plasma; this large difference was also seen previously (Williams et al., 2013).

We identified six groups of five or more miRNAs with similar abundance patterns across biofluids using Bayesian Relevance Network analysis (Ramachandran et al., 2017) (Figure 3, Figure S2). The largest group (group 1) contained 45 miRNAs. A subgroup (1A) containing 31 of these 45 miRNAs had levels that were highest in amniotic fluid and cord blood plasma. All 31 subgroup 1A miRNAs are derived from the 14q32 cluster, which is the largest miRNA cluster in the human genome (54 miRNAs) (Hill et al., 2017). Subgroup 1B contained 14 miRNAs that were highest in cord blood plasma and adult blood plasma and serum but low in amniotic fluid. These include two pairs of miRNAs produced from the same pre-miRNA (miR-486-5p and ‐3p and miR-126-5p and ‐3p) and three miRNAs produced from a cluster on chromosome 17 (miR-451a, miR-144-3p, and miR-4732-3p). Group 2 (23 miRNAs) was enriched for for miRNAs derived from the two miR-200 family clusters, miRs-200b/a/429 and miRs-200c/141. The miR-200 family has important roles in epithelial cells (Korpal and Kang, 2008) and miRNAs in this group were most abundant in seminal plasma and saliva and least abundant in blood-derived biofluids. Group 3 (8 miRNAs) includes 5 miRNAs from the X chromosome miR-506-514 cluster; these miRNAs were highest in ovarian follicle fluid and were also relatively abundant in seminal plasma. Group 4 (23 miRNAs) included a subgroup of 17 miRNAs with highest relative abundance in urine and a second subgroup with 6 members of the miR-34/449 family that were most abundant in CSF, BAL, and amniotic fluid. Group 5 comprises 7 miRNAs that were relatively abundant in bile and Group 6 comprises 5 miRNAs that were relatively abundant in CSF.

**Figure 3.**
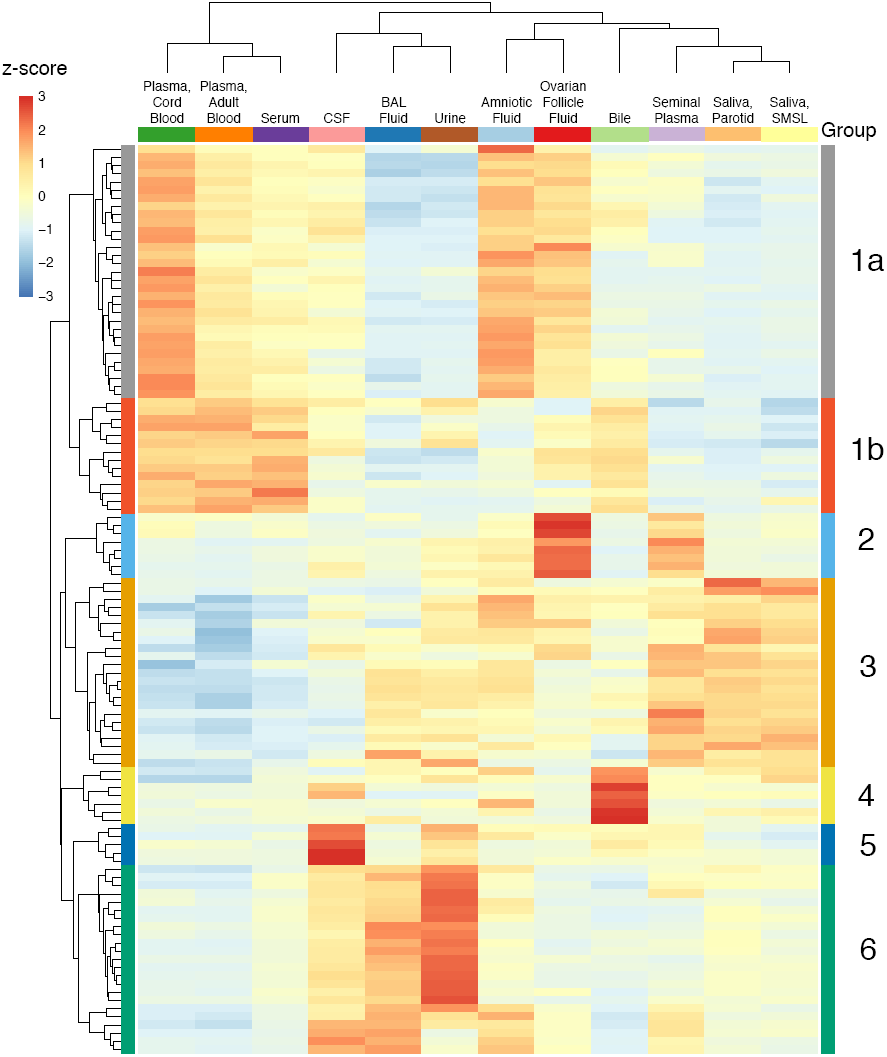
miRNAs with highly correlated read counts across 12 biofluids. Hierarchical clustering heat map depicting scaled miRNA read counts for six groups (1-6) of five or more miRNAs with similar abundance patterns across biofluids using Bayesian Relevance Network analysis. Z-scores indicate levels of miRNA relative to levels of the same miRNA in other biofluids. Figure S2 is a larger version of this figure that includes names of each miRNA.

### tDR profiles

Sequences aligning to tRNAs were detected in all biofluids, although the frequencies of tDR reads varied considerably between biofluids (Fig. 1). Assigning tDRs to the human genome and transcriptome is challenging since many tDRs map to multiple tRNAs corresponding to the same anticodon, a different anticodon for the same amino acid, or even to different amino acids. The Genboree pipeline aggregates tDR reads by amino acid. To analyze tDR reads in more detail, we used MINTmap (Loher et al., 2017), a tDR analysis tool that counts each unique sequence, including those that differ in length by as little as 1 nt, separately. Using this approach, between 954 (bile) and 4,997 (parotid saliva) tDRs had a median of ≥10 reads/million total tDR reads in each biofluid (Fig. 4A, Fig S3, and Table S6). The 10 most frequent tDR reads represent between 18% (adult blood plasma) and 77% (BAL) of total tDR reads (Fig 4B). Pairwise correlation coefficients for tDR read counts between biofluids ranged from 0.08-0.89 (Fig. 4C-E) and were typically far lower than those found for pairwise correlations using miRNA reads. Correlations were highest between adult and cord blood plasma (*R* = 0.89) and between saliva samples from the two different sites (*R* = 0.84). The proportion of mapped reads that represented tDRs was much higher for serum (37%) than for adult plasma (0.8%) and the pairwise correlation for tDR read counts between serum and adult plasma was only moderate (*R* = 0.65). Using all 8,672 tDRs detected in any sample, tSNE analysis revealed that samples of most biofluid types formed largely distinct clusters (Fig. 4E). As with miRNAs, parotid and submandibular/sublingual glands saliva samples formed overlapping clusters. Urine and CSF samples were also overlapping. Although tSNE analysis based on miRNAs produced overlapping clusters of serum and plasma samples, these biofluid types were clearly distinct when the tSNE analysis was based on tDRs.

**Figure 4.**
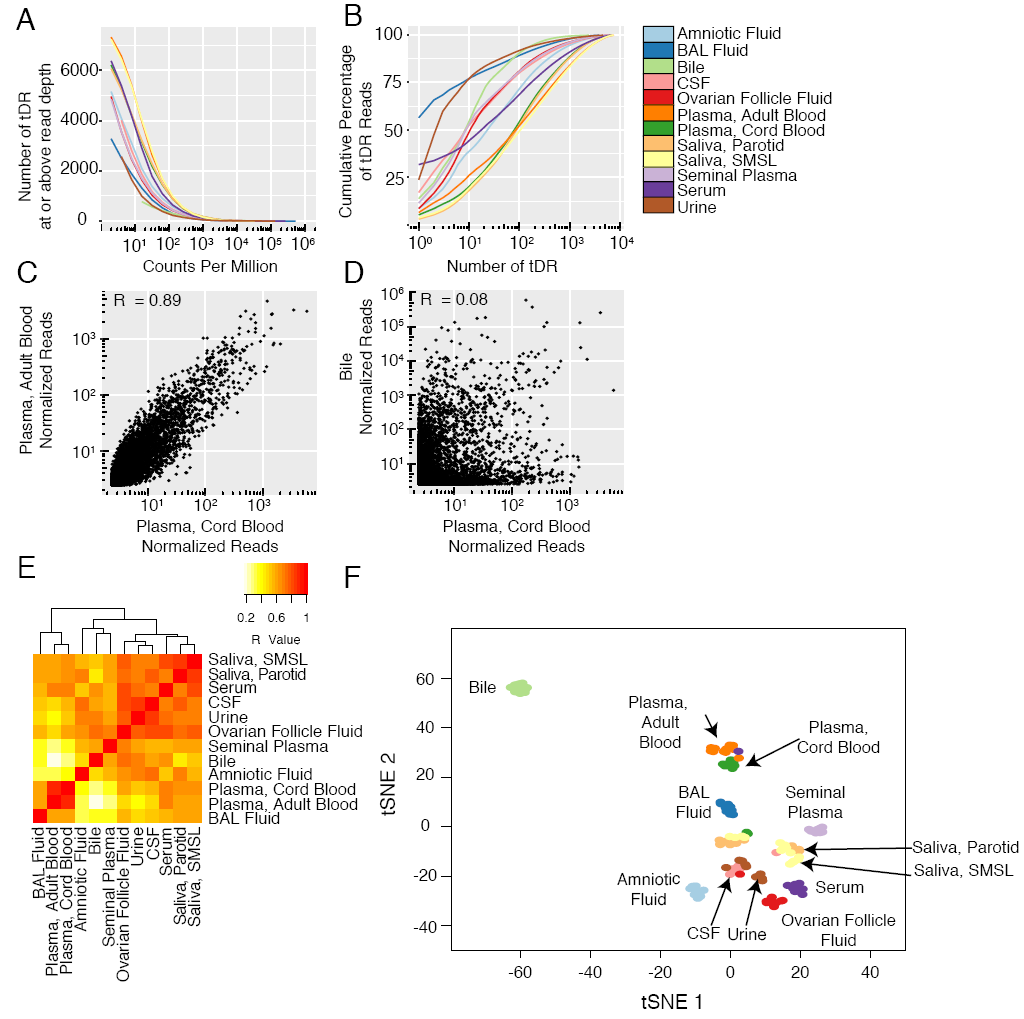
tDR profiles across 12 biofluid types. (A) Number of tDRs detected as a function of read depth. (B) Cumulative distribution of tDR reads. (C and D) Examples of pairwise correlations between biofluids. Each point represents the median normalized read count for a single tDR for the indicated biofluids. One normalized read count was added to each measurement to allow representation of read counts for tDRs with no reads on a log scale. (E) Correlations for all pairs of biofluids. (F) tSNE plot produced using tDR read counts. Each point represents a single biofluid sample.

To further analyze tDRs, we grouped together tDRs based on the amino acid that corresponds to the tRNA of origin (Fig. 5A and Fig. S4). tDRs that could not be unambiguously assigned to an amino acid were excluded from this analysis. Glycine tDRs were predominant in urine, serum, and CSF. Leucine tDRs were predominant in BAL (62%) but were much less frequently found (<15%) in other biofluids. Glutamic acid tDRs were predominant in amniotic fluid, bile, plasma from cord blood and adult blood, and both types of saliva, whereas methionine tDRs were predominant in seminal plasma. Glycine, glutamate, and methionine each represented >20% of tDR reads in ovarian follicle fluid. Median tDR reads for tyrosine, asparagine, phenylalanine, and isoleucine tDRs were <1% each for all biofluid types. Normalized read counts for many tDR-associated amino acids differed markedly between biofluids. For example, tRNA reads aligning to glycine made up 68.9% of tDR reads in urine but ≤43.4% of tDR reads in every other biofluid. A short (16 nt) read mapping to the 5’ end of leucine tRNAs accounted for the majority (59%) of all tDR reads in BAL fluid samples but was rare in all other biofluids (0.003%-1.6%).

**Figure 5.**
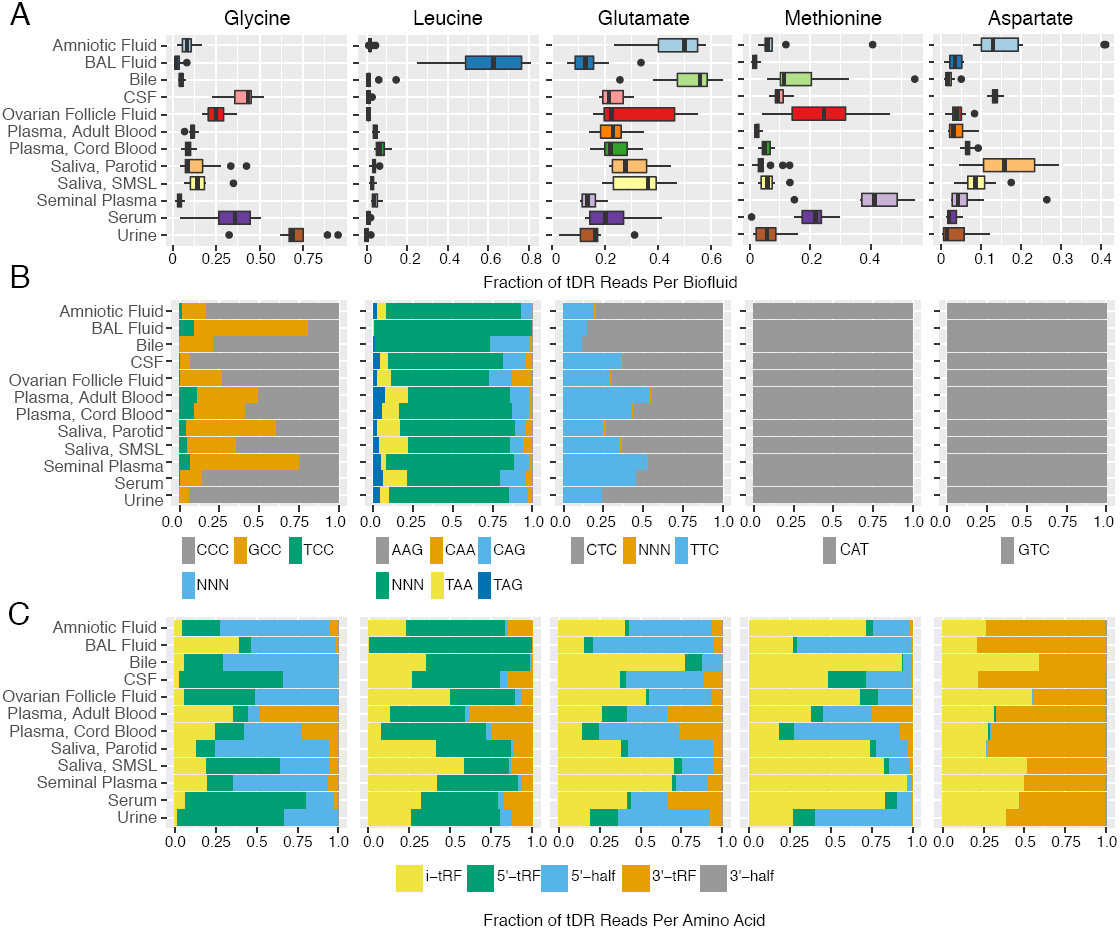
tDR abundance by amino acid, anticodon, and fragment type. (A) tDR abundance by amino acid. (B) tDR abundance by anticodon. (C) tDR abundance by fragment type. Data are shown for tDRs from the five most highly represented tRNAs. Data for other tDRs are shown in Figures S4-S6.

For amino acids encoded by more than one codon, we examined the distribution of reads for each possible anticodon (Fig. 5B and Fig. S5). For some amino acids (e.g., glycine), anticodon frequency varied substantially between biofluids but for others (e.g., leucine) anticodon frequency was less variable. We also classified tDRs by fragment type (Fig. 5C and Fig. S6). Fragment types differed substantially according to the tRNA of origin. For example, in most biofluids glycine tDRs mapped primarily to the 5’ region (5’ tRFs and 3’ halves), whereas methionine and histidine tDRs mapped primarily to internal regions of tRNAs (i-tRF). To examine differences in tDR fragments between biofluids in more detail, we constructed read coverage maps for four tDRs with the largest numbers of mapped reads (Fig. S7). Inspection of these coverage maps reveals large differences in fragment types across biofluids (e.g., increased 3’ coverage in amniotic fluid for three of these four tRNAs) and illustrates major differences in fragment sizes (e.g., shorter 5’ fragments for tRNA7 LeuAAG compared with the other three tRNAs).

### Y RNA profiles

We found reads mapping to each of the four human Y RNA genes in every biofluid (Figure 6A and Tables S7 and S8). In each biofluid except for BAL fluid, most reads that mapped to Y RNA mapped to RNY4. In BAL fluid, the mean mapping rate for RNY4 was 43% and RNY5 accounted for 49%. Most reads mapped to either the 5’ region or the 3’ region of full-length Y RNAs, with relatively few reads mapping to the central portion of Y RNAs (Figure 6B). The proportion of 5’ reads to 3’ reads varied between biofluids and between different Y RNA genes. For example, RNY4 fragments more frequently mapped to the 5’ end in seminal plasma and CSF but more frequently mapped to the 3’ end in adult blood plasma, BAL fluid, and saliva.

**Figure 6.**
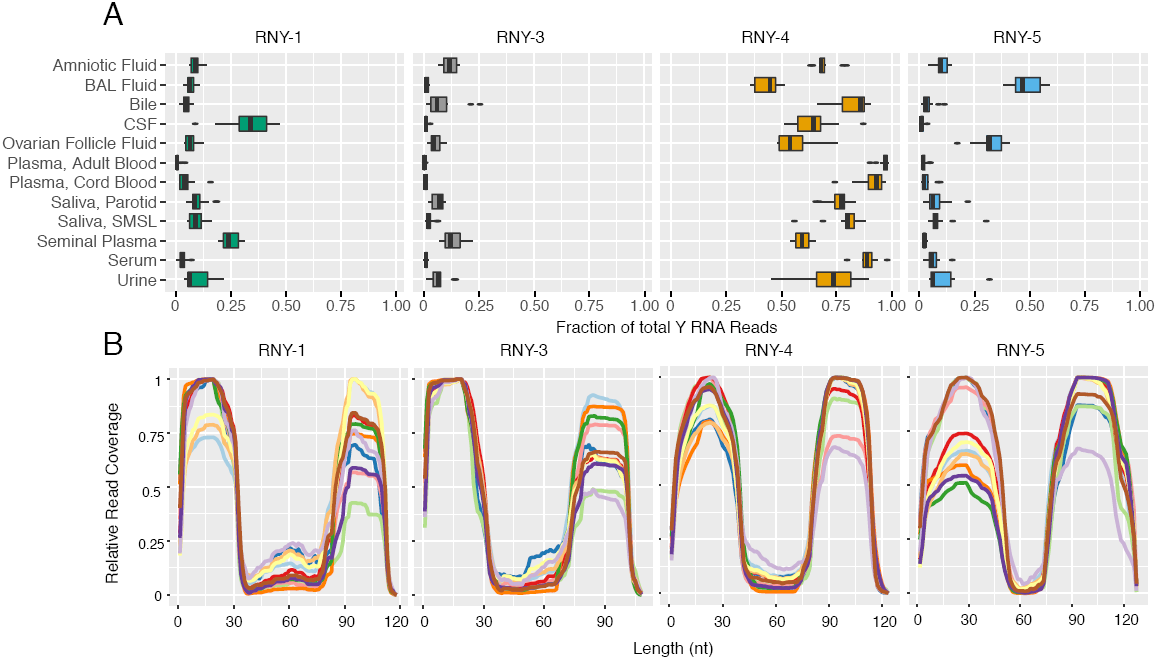
Y RNA fragments in 12 biofluid types. (A) Distribution of Y RNA reads by Y RNA gene. (B) Y RNA read mapping positions. We determined the number of reads covering each nucleotide of each full-length Y RNA. Values are normalized to the position of each Y RNA with the largest number of reads in each biofluid.

### piRNA profiles

As discussed previously, some biofluids contained substantial proportions of reads that mapped to both piRNAs and other Gencode RNAs (especially RNY4). The majority of reads that mapped to piRNAs also mapped to other RNA biotypes except in one biofluid, seminal plasma. 57% of seminal plasma reads that mapped to piRNAs did not map to another biotype. After exclusion of ambiguously mapped reads (see Experimental Procedures), the number of piRNAs detected was higher in seminal fluid (5588 distinct piRNAs in one or more samples) than in other fluids (94-1068 piRNAs). The most frequent 57 piRNAs accounted for 50% of seminal plasma reads that mapped exclusively to piRNAs (Table S9).

## DISCUSSION

We applied a standardized RNA-seq approach to identify small RNAs in 12 human biofluid types. This work extends previous analyses of biofluid extracellular RNAs by applying a method that allows for relative quantification of small RNAs from multiple biotypes to a diverse collection of normal biofluids. We found small RNAs that mapped to transcripts of many biotypes. In each biofluid tested, the majority of mapped reads could be assigned to miRNAs, tDRs, or Y RNAs. Additional reads mapping to piRNAs, mRNAs, long non-coding RNAs, small nuclear RNAs, and other RNAs were also detected but were less common. A major finding was that the relative levels of different RNA biotypes differed dramatically between biofluids. The relative abundance of the two major biotypes, miRNAs and tDRs, varied by >10^3^-fold across the set of 12 biofluids. Our results are consistent with those from another recent study that found relatively high levels of tDRs in urine and relatively high levels of Y RNA fragments in plasma (Yeri et al., 2017). Despite differences in overall small RNA composition, miRNA levels were generally highly correlated between biofluids (*R* = 0.79-0.98). In contrast, tDR levels were less well correlated (*R =* 0.08-0.89). Biofluids that had highly similar profiles for small RNAs of one biotype sometimes had very different profiles for RNAs of other biotypes. For example, adult blood plasma and serum miRNA levels were highly correlated (*R* = 0.98) but tDR levels in these biofluids were much less well correlated (*R* = 0.68). Our findings demonstrate the extensive diversity of small RNA populations within each biofluid type and highlight both similarities and differences between biofluids.

The methods we used have advantages and limitations compared with those used in previous studies. A major feature of our work is that we used a uniform approach to RNA isolation, RNA-seq library preparation, sequencing library size selection, sequencing, and data analysis for all samples. Like some other studies, we used RNA-seq rather than other methods such as PCR‐ or hybridization-based methods that focus only on certain predefined RNAs. The RNA-seq method we used has been shown to reduce bias by employing oligonucleotides with random nucleotide sequences for adaptor ligation (Giraldez et al., 2017) and we used a permissive size selection strategy rather than one designed to enrich for small RNAs in the miRNA size range. We used a publicly available analysis pipeline and our data are publicly available, which allows for re-analysis of the primary data as databases and analysis tools are updated. To obtain a broad view of small RNAs in biofluids, we extracted RNA from unfractionated biofluids. Other studies have shown differences in small RNA content of specific biofluid compartments such as exosomes and argonaute 2-containing ribonucleoprotein complexes (Arroyo et al., 2011), and further work will be required to understand the compartmentalization of RNAs in the large set of biofluids we studied. While our method detected many RNAs belonging to multiple biotypes, the method is designed for sequencing of RNAs with a 5’ phosphate and a 3’ hydroxyl group and not for RNAs with other end modifications or post-translational modifications such as those frequently found in tRNAs. Finally, the method we used is well suited for relative quantification of each RNA species across multiple samples, but different approaches will be necessary for rigorous measurements of absolute quantities of RNAs in different biofluids.

The miRNA profiles of these 12 biofluids were remarkably similar overall. Four miRNAs were among the top 10 detected in most biofluids (miR-92a-3p in 10 biofluids, miR-99a-5p in 9 biofluids, and miR-24-3p and miR-26a-5p in 7 biofluids; see Table S3). A previous report also identified a small group of miRNAs as being highly abundant in many biofluids (Weber et al., 2010). Seven biofluid types from that prior PCR-based study (adult blood plasma, urine, seminal plasma, saliva, amniotic fluid, BAL, and CSF) were included in the set of biofluids we analyzed by small RNA-seq. The prior study identified nine miRNAs that were ranked within the top 10 miRNAs in at least 4 of these 7 biofluids: miR-335* (now known as miR-335-3p), miR-515-3p, miR-518e, miR-616* (miR-616-5p), miR-302d, miR-892a, and miR-509-5p. None of those miRNAs were identified as being among the top 10 miRNAs in any of the biofluids in our RNA-seq analysis. This likely reflects differences in specificity or bias between the methods used in the two studies. A recent evaluation of the RNA-seq-based method we used showed good performance characteristics for analysis of human extracellular miRNAs and low bias as assessed using large panels of synthetic miRNAs (Giraldez et al., 2017). Nonetheless, RNA-seq and other methods in common use (including PCR-and hybridization-based methods) are better suited for comparing levels of each miRNA between samples rather than comparing levels of different miRNAs within each sample.

We identified large differences in levels of some miRNAs between biofluids. These may arise from differences in local production of cellular miRNAs. Consistent with that idea, we found higher levels of extraembryonic cell miRNAs in amniotic fluid, nervous system cell miRNAs in CSF, and an epididymal miRNA in seminal plasma. We found many cases in which miRNAs produced from the same pre-miRNA or the same miRNA cluster had highly correlated levels across the full set of miRNAs, which also suggests that miRNA production is a major determinant of differences in miRNA abundance. In addition to differences in local production of cellular miRNAs, differences in extracellular miRNAs could arise from preferential secretion of miRNAs from cells (Valadi et al., 2007) and from differences in processing or elimination of extracellular miRNAs. Since extracellular miRNAs have been shown to be capable of entering cells may modulate gene expression (Patton et al., 2015), our results suggest that differences in miRNA composition between biofluids may be functionally important.

We found large differences in tDRs across biofluids. tRNAs can be processed into tDRs by specific enzymatic cleavage events and tDRs have been previously identified in human biofluids, including human serum and urine, and in exosomes from human semen (Dhahbi, 2015; Yeri et al., 2017). We classified tDRs according to the tRNA of origin (amino acid and anticodon) and the fragment type (5’ or 3’ halves or 5’, 3’, or intermediate fragments) and found differences in each of these classifications across the set of biofluids. Many factors, including tRNA levels, tRNA cleavage, tDR secretion, and biodistribution and elimination of extracellular tDRs may contribute to determining the levels of tDRs in each biofluid. Intracellular tDRs have been reported to have many roles, including inhibiting protein translation during cellular stress (Ivanov et al., 2011) and mediating intergenerational inheritance (Chen et al., 2016; Sharma et al., 2016). The roles of extracellular tDRs remains enigmatic, but the presence of relatively high levels of extracellular tDRs in biofluids such as bile, urine, seminal plasma, and amniotic fluid (where tDRs represented the majority of all mapped small RNA reads) suggests that extracellular tDRs may be functionally important. At least some of the functional roles of intracellular tDRs are quite specific: for example, a tDR from the Gly-GCC tRNA but not other tDRs were shown to regulate retroelement-driven transcription of genes that are highly expressed in preimplanation embryos (Sharma et al., 2016). Differences in tDR populations between biofluids may therefore have important functional implications.

Y RNA fragments represented a substantial proportion of the small RNAs that we detected in some biofluids, including adult and cord blood plasma, serum, and CSF. A previous RNA-seq study also reported that Y RNA fragments made up a large proportion of small RNA reads in human serum and plasma (Dhahbi et al., 2013). That study found that >95% of fragments mapped to the 5’ end of Y RNAs. Our results, obtained using an RNA-seq method developed to reduce bias, revealed a substantially higher proportion of 3’ Y RNA fragments. Intact cellular Y RNAs are required for the initiation of DNA replication, regulation of RNA stability, and cellular responses to stress (Kowalski and Krude, 2015). The role of Y RNA fragments is less well understood. It has been reported that RNY5 fragments produced in cancer cell extracellular vesicles trigger rapid cell death in primary cells but not in cancer cells (Chakrabortty et al., 2015), suggesting that further studies of the biogenesis, trafficking, and function of Y RNA fragments found in biofluids are warranted.

We found that blood plasma and serum contained a substantial number of small RNAs with sequences that map to piRNAs. A previous study also detected many sequences that mapped to piRNAs in human plasma (Freedman et al., 2016). However, we urge caution about inferring that small RNA reads that map to piRNAs are derived from piRNAs. The large majority of piRNA-mapped reads we obtained in our plasma and serum samples also mapped to other RNA biotypes (especially Y RNAs) that are abundant in these biofluids. In our study, the piRNA with the largest number of mappings shares sequence with a fragment of RNY4, a Y RNA. The most frequent piRNA-mapped read in the prior study (Freedman et al., 2016) was piR-33043 (Genbank accession DQ592931), which also shares sequence with RNY4. We therefore confined our analysis of piRNAs to sequences that did not map to RNAs of other biotypes. piRNAs have prominent roles in the testis (Girard et al., 2006) and even when using this more conservative approach we found that a substantial number and variety of piRNA sequences were detected in seminal plasma. A previous report also identified piRNAs in seminal plasma and identified a set of piRNAs that were associated with infertility (Hong et al., 2016). Although previously suggested to be germline-specific, piRNAs also play roles in non-gonadal cells (Ishizu et al., 2012) and non-gonadal cells might therefore be a source of piRNAs in biofluids outside the reproductive tract. However, we found relatively few reads that mapped only to piRNAs in biofluids other than seminal plasma. The functional significance of extracellular piRNAs is not yet known.

By using a uniform RNA-seq-based approach, we produced an extensive catalog of small RNAs that are present in a large set of human biofluids. The RNA composition of biofluids differed widely, with major differences in the distribution of biotypes and of specific RNAs within each biotype. This work provides a resource for investigators seeking to understand the production, distribution, and function of extracellular RNAs. Biofluid small RNAs are being explored as disease biomarkers and our work also helps to identify RNAs that are present in multiple biofluids and may have potential as novel biomarkers.

## ACKNOWLEDGEMENTS

We thank each of the individuals who participated in the study by providing samples. We thank Christine P. Nguyen for assistance with collection and annotation of the BAL fluid, adult blood plasma, serum, and urine samples. We thank Serena Spudich for access to CSF samples collected as part of studies that she directed. This work was supported by National Institutes of Health Extracellular RNA Communication Program (grants U01HL126493, to P.G.W. and D.J.E. and UH2TR000884 and UH3TR000884, to T.P.).

## AUTHOR CONTRIBUTIONS

P.M.G., N.R.B., A.J.B., P.G.W., and D.J.E. designed the study. N.R.B., H.C., S.F., T.C.M., T.P, R.W.P., J.F.S., and P.G.W. recruited study participants and collected biofluid samples. P.M.G. performed the RNA-seq experiments and analyzed the data. P.M.G. and D.J.E. wrote the paper. All authors contributed to manuscript editing.

## DECLARATION OF INTERESTS

The authors declare no competing interests.

## EXPERIMENTAL PROCEDURES

### Human Subjects

With the exception of bile samples, which were obtained at the Mayo Clinic following liver transplantation, all samples were obtained from healthy subjects enrolled in a variety of studies conducted at the University of California, San Francisco. Protocols were approved by Institutional Review Boards at the Mayo Clinic (bile) and the University of California San Francisco (all other specimen types). Except for cord blood plasma, which was obtained at birth, all other samples were obtained from studies enrolling subjects aged 18 years or older. Details of sample collection and storage for each sample type are provided in Supplemental Experimental Procedures.

### RNA Isolation, Library Preparation, and Sequencing

After thawing, samples were spun at 2,000 x *g* for 5 minutes at 4°C. RNA was isolated from 200 μL of biofluid using the Qiagen miRNEasy Micro Kit according to the manufacturer’s protocol except that 1 mL of Qiazol and 180 μL of chloroform were used. Small RNA-seq libraries were prepared using 4N protocol D as previously described (Giraldez et al., 2017). This method relies on a version of the TruSeq Small RNA Library Preparation Kit modified by using randomized adapters, adding PEG to the adapter-RNA solution, and including steps to enzymatically remove excess adapter after 3’ ligation. Multiple libraries with unique indexes were pooled, purified using the Qiagen MinElute PCR Purification Kit per the manufacturer’s recommendations. Libraries were size-selected using the PippinPrep (Sage Science) with a 3% agarose gel. In pilot experiments, we adjusted the size selection parameters to maintain a low proportion of adapter dimers (132 bp) and maximize the proportion of library with inserts of ∼22-30 bp. We selected the PippinPrep broad range option to deplete sequences <137 bp and >166 bp but maintain a range of insert sizes within this range. Size-selected DNA was sequenced on an Illumina HiSeq 4000 (single end 50 base mode).

### Sequence alignment

All FASTQ files were processed using the exceRpt small RNA-seq pipeline version 4.6.2 available on the Genboree Workbench (http://www.genboree.org). Sequence reads were clipped 4 nt from the invariant portions of both 5’ and 3’ adapters to remove the 4 randomized nt in the adapters. Clipped reads were aligned with a one mismatch allowance. To exclude piRNAs that mapped to other biotypes, we changed the order of mapping assignment to count Gencode alignments in preference to piRNA alignments (alignment order: miRNAs, tRNAs, all annotations from Gencode, piRNAs, and circular RNA). We also eliminated piRNA mappings if the large majority (>98%) of reads mapping to a given piRNA also mapped to an RNA of a different biotype.

### RNA-seq data inclusion criteria

RNA-seq results were included if: 1) ≥100,000 reads mapped to the transcriptome, 2) the number of reads mapping to annotated transcripts represented at least 50% of the number of reads mapping to the human genome, and 3) *R* ≥ 0.85 for correlation of miRNA reads with most other samples of the same biofluid. For samples that failed to meet inclusion criteria on initial analysis, duplicate biofluid samples were used for a second round of RNA isolation and library preparation. If the repeat analysis met inclusion criteria, these results were used for all analyses. If the repeat analysis did not meet the inclusion criteria, data from that sample were not used for subsequent analyses. There was no case in which replicate analysis of a sample gave consistent results (*R* ≥ 0.85 for miRNAs) but these results were poorly correlated (*R* < 0.85) with most other samples of the same biofluid type.

### RNA Biotype Distribution Analysis

We used the biotype counts file generated by the small RNA-seq pipeline on the Genboree Workbench. Since Y RNA reads represented a large proportion of Gencode transcript reads in some biofluids, we used the Gencode read counts file to quantify Y RNA reads separately from other Gencode reads.

### miRNA analysis

miRNA diversity analyses were performed using the mean normalized miRNA read counts for all samples of a given biofluid type. Read counts for each miRNA in each sample were normalized by dividing by the total number of miRNA read counts in that sample. For analyses of the numbers of miRNAs detected as a function of read depth, we excluded all miRNAs with mean read counts < 1/10^6^ total miRNA reads (lower limit of detection). For biofluids where miRNAs represented a small proportion of total RNA reads, the total number of miRNA reads for all samples was <10^6^. For those biofluids, the lower limit of detection was defined as 10^6^ divided by the total number of miRNA reads for that biofluid. For analyses of cumulative distributions, we ranked miRNAs in descending order of normalized mean read counts for a given biofluid and calculated the cumulative sum until we included enough miRNAs to account for 99.9% of miRNA reads.

For pairwise correlations and tSNE analyses, we normalized read counts by DESeq2 (Love et al., 2014) and determined mean read counts for all samples of each biofluid. We calculated the Pearson correlation coefficient of each biofluid pair using R. We generate tSNE plots using the Rtsne function of the Rtsne R package with perplexity 20 and a maximum iteration of 5,000. We plotted the results using the xyplot function from the lattice R package. We used Bayesian Relevance Networks (Ramachandran et al., 2017) to generate a list of co-expressing miRNAs. We normalized miRNA read counts with DESeq2, transformed the counts data using the varianceStabilizingTransformation function, identified the top 25% most frequent miRNAs (by mean), and then selected the top 13% most variable as input to the Bayesian Relevance Networks algorithm. We selected all co-expressing miRNAs with a Bayesian correlation ≥ 0.80, which had an estimated false discovery rate of 0.012. We used the pheatmap function from the pheatmap R package to generate a hierarchical clustering diagram for all miRNAs belonging to networks with 3 or more miRNAs.

### tDR analysis

We used MINTmap (Loher et al., 2017) to analyze tDRs. We removed adapters, trimmed 4 nucleotides from each end of the read, and removed low quality reads (using standard Genboree parameters) from all fastq files with the fastx-toolkit. Fastq files were then processed individually by MINTmap. We only counted alignments that mapped exclusively to annotated tRNA regions to reduce ambiguity. For analyses of amino acid or anticodon read counts, reads that mapped ambiguously (to multiple amino acids or anticodons) were deemed “undetermined.” All reads were aggregated by sum and normalized by the total number of tDR reads within each sample or biofluid type. To analyze tRNA coverage, we took the normalized mean of each sample and aggregated read counts by biofluid and the start and end position.

### Y-RNA analysis

Using the bam files generated by the Genboree pipeline using the “Upload Full Results” option, we searched for any alignment to the 4 human Y RNAs (RNY1-201, RNY3-201, RNY4-201, and RNY5-201) and considered each unique fragment a distinct read. To plot read coverage, we followed the same strategy as outlined above for tDRs.

### Data Availability

Data are available through the exRNA Atlas (https://exrna-atlas.org) and as supplemental tables. For two biofluid types (parotid saliva and SMSL saliva), the IRB-approved consent process permitted release of read counts but not sequences and is available through accession number EXR-DERLE1PHASE1PROT-AN. For the other 10 biofluid types, both raw sequence data and read counts are available (EXR-DERLE1PHASE1OPEN-AN).

